# Genome Report: First whole genome assembly of *Python regius* (ball python), a model of extreme physiological and metabolic plasticity

**DOI:** 10.1101/2025.02.01.635752

**Authors:** Dakota R. Hunt, Holly Allen, Thomas G. Martin, Sophia N. Feghali, Edward B. Chuong, Leslie A. Leinwand

## Abstract

The study of nontraditional model organisms, particularly those exhibiting extreme phenotypes, offers unique insights into adaptive mechanisms of stress response and survival. Snakes, with their remarkable physiological, metabolic, and morphological adaptations, serve as powerful models for investigating these processes. Pythons are a unique model organism that have been studied for their extreme metabolic and physiological plasticity. To date, the Burmese python (*Python bivittatus*) is the only member of the *Pythonidae* family to have been sequenced. The low contiguity of this genome and rising challenges in obtaining Burmese pythons for study prompted us to sequence, assemble, and annotate the genome of the closely related ball python (*Python regius*). Using a hybrid sequencing approach, we generated a 1.45 Gb genome assembly with a contig N50 greater than 18 Mb and a BUSCO score of 98%, representing the highest quality genome to date for a member of the *Pythonidae* family. This assembly provides a valuable resource for studying snake-specific traits and evolutionary biology. Furthermore, it enables exploration of the molecular mechanisms underlying the remarkable cardiac and muscular adaptations in pythons, such as their ability to rapidly remodel organs following feeding and resist muscular atrophy during prolonged fasting. These insights have potential applications in human health, particularly in the development of therapies targeting cardiac hypertrophy and muscular atrophy.

## Introduction

The study of nontraditional model organisms displaying extreme phenotypes can facilitate the discovery of adaptive mechanisms of stress response and survival. Such mechanisms, which can be difficult to identify or not present in common model organisms, may provide novel insight into human biology and drive innovative therapeutic approaches to human disease (Stenvinkel et al. 2020). Snakes represent one such model due to their significant metabolic, physiological, and morphological adaptations acquired by evolving in environments that would be considered extreme for mammals (Castoe et al. 2008; Vonk and Richardson 2008). Pythons, in particular, have been studied as a vertebrate model of extreme metabolic and organ structural plasticity due to their consumption of large prey after months of fasting (Secor and Diamond 1995; Secor and Diamond 1998). The post-prandial metabolic and physiological remodeling of digesting pythons is unparalleled in known biology in both magnitude and rapidity (Tan et al. 2023; Martin et al. 2024). Pythons display significant organ remodeling, metabolic alterations, and associated changes in gene expression and chromatin remodeling in response to feeding (Secor and Diamond 1995; Castoe et al. 2008; Riquelme et al. 2011; Crocini et al. 2024; Martin et al. 2024). For example, python hearts undergo up to a 40% increase in mass 2-3 days after consuming a large meal, followed by regression to the just above their original size within the following days (Secor and Diamond 1995; Secor and Diamond 1998; Andersen et al. 2005; Riquelme et al. 2011). Intriguingly, pythons do not appear to lose muscle mass during these long fasting periods (McCue 2007), nor do their hearts undergo atrophy beyond a baseline over prolonged periods of fasting (Martin et al. 2024).

Currently, the Burmese python (*Python bivittatus*) is the only member of the *Pythonidae* family to have its genome sequenced (Castoe et al. 2013). The low quality/contiguity of this genome can be attributed to the usage of shotgun sequencing approaches. Since then, the advent of long-read sequencing technologies, the reduced cost of short-read sequencing, and improvements in genome assembly tools have resulted in more contiguous and better annotated genomes (Whibley et al. 2021). While most studies on python biology have utilized Burmese pythons, recent work has included the use of Ball pythons (*Python regius*, Crocini et al. 2024). Furthermore, challenges in sourcing Burmese pythons for study, brought on by their invasive species designation in the United States, has led our laboratory and others to employ Ball pythons as a model organism (Fig. 1).

**Figure 1.**
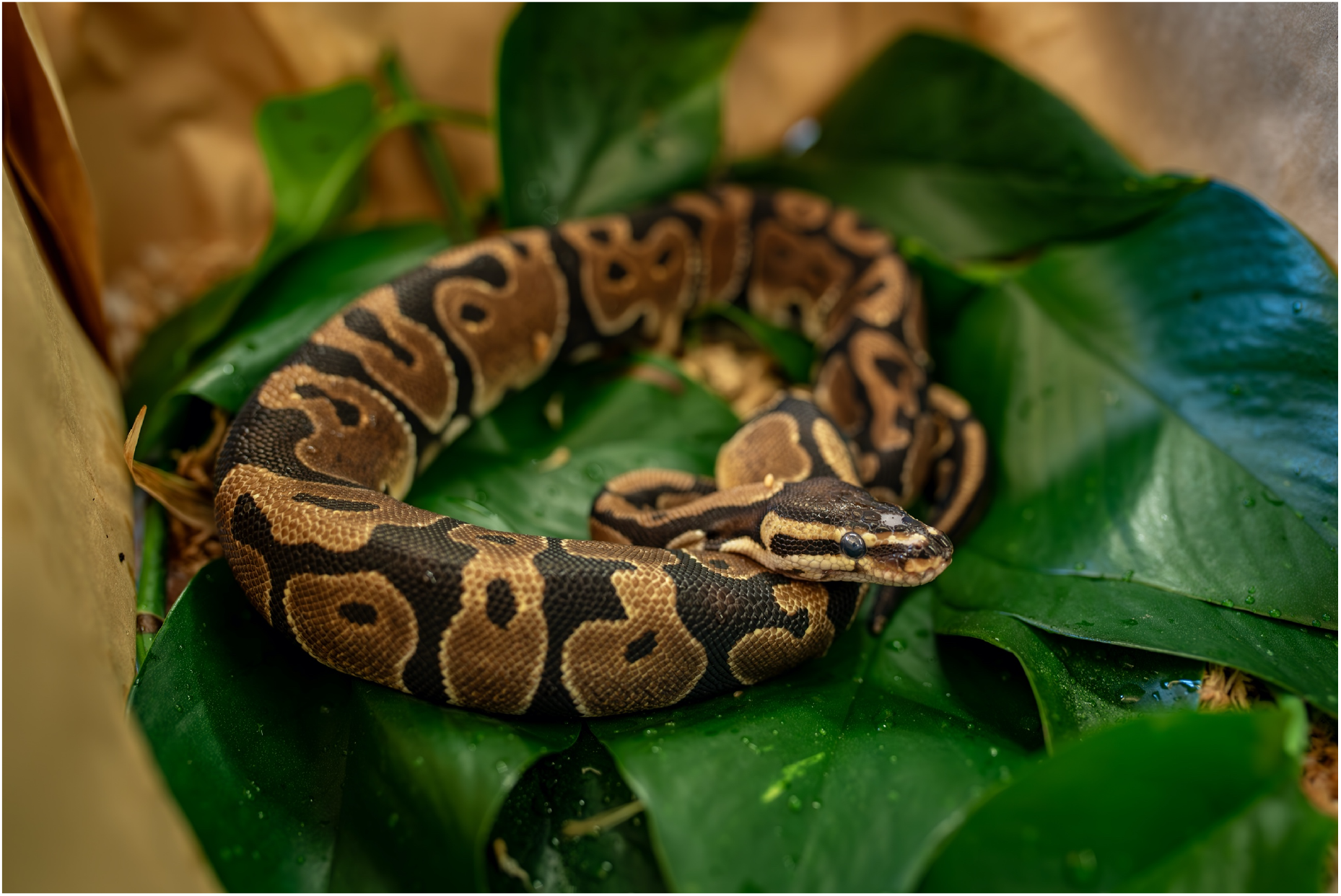
Representative image of *Python regius*. Photo taken by Yuxiao Tan.

Here we present the first *P. regius* genome assembly, and the first high quality genome (vertebrate benchmarking universal single-copy ortholog [BUSCO] scores > 90%, contig N50 > 1 Mb) for a member of the *Pythonidae* family. This resource will not only expand our understanding of snake biology and evolution of snake-specific traits, but also aid in future studies of unique physiological adaptations, particularly in the realms of cardiac biology and resistance to muscular atrophy. Such work may directly contribute to novel therapeutic approaches targeting pathological cardiac hypertrophy or muscular atrophy in humans.

## Methods & Materials

### Specimen origin and tissue collection

All animal procedures were approved by the University of Colorado Boulder Institutional Care and Use Committee (IACUC). The ball python (*Python regius*) specimen used in this study was a captive-bred adult female received from Bob Clark Reptiles (Oklahoma City, OK, USA). The specimen was housed at the University of Colorado Boulder BioFrontiers Institute vivarium in a room kept at 28°C and 50% relative humidity. The housing room was subjected to cycling 12-hour light-dark periods. This specimen was fasted for 28 days before being fed a rat meal equivalent to 25% of its body weight. The specimen was then fasted for 70 days before being anesthetized with isoflurane and subsequently euthanized via rapid decapitation. The specimen was dissected and tissues (cardiac ventricle, liver, skeletal muscle, small intestine, brain, kidney, cardiac adipose, visceral adipose, ovary) were collected and snap frozen in liquid nitrogen before being stored at −80°C.

### DNA extraction and sequencing

High molecular weight (HMW) genomic DNA (gDNA) was extracted and purified using the Monarch^®^ HMW DNA Extraction Kit for Tissue from New England Biolabs (Ipswich, MA, USA). 15 mg of frozen small intestine was biopulverized and further homogenized using the provided pestle. Manufacturer’s instructions were followed to extract and purify the DNA. Samples were lysed in a thermal mixer at 56°C, 1400 rpm for 45 minutes. Eluted HMW gDNA was incubated overnight at room temperature to ensure homogenous dissolution. Quantity and quality of isolated gDNA was verified by NanoDrop and an Agilent 2200 TapeStation (Agilent Technologies, Santa Clara, CA, USA) was used to verify the gDNA was indeed of a high molecular weight. A second, standard, genomic DNA extraction was performed to generate additional Nanopore reads. DNA was extracted from ∼50 mg of small intestine using the Quick-DNA Tissue/Insect Kit (Zymo Research), according to the manufacturer’s instructions.

Genomic libraries were prepared from both gDNA preps with the Native Barcoding Kit 24 V14 kit (Oxford Nanopore Technologies, Oxford) and sequenced on a R10.4.1 PromethION flow cell (Oxford Nanopore Technologies). HMW gDNA was sent to the Genomics Shared Resource Core at the University of Colorado Anschutz Medical Campus (Aurora, CO, USA) for short-read whole genome sequencing. Library preparation was performed using the Ovation® Ultralow System V2 library preparation kit (Tecan, Männedorf, Switzerland). Whole genome sequencing was performed using the Illumina NovaSeq X platform (Illumina, San Diego, CA, USA) at a requested coverage of 60x.

### Genome assembly and polishing

Dorado v0.8.2 (Oxford Nanopore Technologies, Oxford, United Kingdom) was used to basecall reads from raw Nanopore sequencing data via the super high accuracy model. Dorado was then used to demultiplex reads with the --no-classify and --emit-fastq flags. NanoStat (De Coster et al. 2018) was used to generate read statistics. *De novo* assembly of the ball python genome was conducted using Flye v2.9.5 (Kolmogorov et al. 2019) with an estimated genome size of 1.5 Gb (--genome-size 1.5g). Medaka v2.0.1 (Oxford Nanopore Technologies, Oxford, United Kingdom) was used to polish the draft Flye assembly using the raw basecalled Nanopore reads and the r1041_e82_400bps_sup_v5.0.0 model. The bbduk.sh script from BBMap v38.05 (Bushnell) was used to trim adapters from the sequenced Illumina short reads with the following parameters: ktrim=r k=31 mink=11 hdist=1 tpe tbo qtrim=r trimq=10. Trimmed reads were then aligned to the Medaka-polished genome by BWA-MEM v0.7.15 (Li 2013). Samtools v.1.16.1 (Li et al. 2009; Danecek et al. 2021) was used to create a sorted and indexed bam file containing the aligned reads. Low quality (-q 10) and unmapped (-F 4) reads were filtered out prior to sorting. Pilon v1.24 (Walker et al. 2014) was used to further diploid-aware polish the genome assembly using the aligned Illumina short reads.

### Genome quality control

The Quality Assessment Tool (QUAST) v5.3.0 (Mikheenko et al. 2023) was used to generate genome metrics (total length, N50, L50, etc.) and assess genome quality. Benchmarking Universal Single Copy Orthologs (BUSCO) v5.8.2 (Manni et al. 2021) is a tool used to assess genome assembly quality and completeness by identifying the presence and integrity of a set of universally conserved single-copy orthologous genes across species. The polished Pilon assembly was input into BUSCO and run against the following lineages: ‘sauropsida_odb10’ (7480 BUSCOs) and ‘vertebrata_odb10’ (3354 BUSCOs).

### Assembly decontamination and filtering with Blobtoolkit

Blobtoolkit v4.4.0 (Challis et al. 2020) was used to perform further QC and screen for potential contamination within the genome assembly. Blobtoolkit integrates genome coverage information, BLAST results, and BUSCO analysis output to allow for interactive visualization of a dataset and subsequent filtering of an assembly dataset. To generate coverage information, we aligned the Illumina and Nanopore reads back to the Pilon-polished genome. Illumina reads were aligned using BWA-MEM and basecalled, demultiplexed Nanopore reads were aligned using minimap2 v2.17 (Li 2018). Samtools was used to convert the subsequent sam files to bam files, and sort and index the bam files. To generate taxonomic information for each contig in the Pilon-polished assembly, we used the Basic Local Alignment Search Tool (BLASTn 2.7.1+) (Altschul et al. 1990; Camacho et al. 2009). Assembly contigs were queried against the core_nt database v1.1 from the National Center for Biotechnology Information (NCBI) using the following flags: *-outfmt “6 qseqid staxids bitscore std” -max_target_seqs 10 -max_hsps 1 -evalue 1e-25*. DIAMOND v2.1.10 (Buchfink et al. 2021) is a protein and translated sequence aligner that is significantly faster than BLAST. Diamond makedb was used to create a Diamond-formatted database from the curated, non-redundant UniProtKB/Swiss-Prot protein sequence database. Taxonomic information was added using the “prot.accession2taxid.FULL”, “names.dmp”, and “nodes.dmp” files from NCBI. Diamond was then used to blast the query Pilon-polished assembly against the created database with the following flags: *--outfmt 6 qseqid staxids bitscore qseqid sseqid pident length mismatch gapopen qstart qend sstart send evalue bitscore --sensitive --max-target-seqs 1 --evalue 1e-25*. A Blobtools dataset was created using the Blobtools *create* command and specified taxonomic information for *Python regius*. The *add* command was then used to add coverage information, BLASTn/DIAMOND hits, and BUSCO output to the dataset. The Blobtools *filter* command was used to remove all contigs less than 3000 base pairs from the assembly.

### Purge_dups

Purge_dups v1.0 (Guan et al. 2020) was used to remove low-coverage contigs, collapsed repeat contigs, haplotigs, and contig overlaps from the Blobtools-filtered genome assembly. A configuration file specifying the input Nanopore and Illumina data was generated using the pd_config.py script and the pipeline was run using the run_purge_dups.py script. The high coverage cutoff was then manually changed, resulting in the following coverage cutoffs: low=5, mid=21, high=100. The Blobtools-filtered assembly was then purged using these manual cutoffs and the -c and -e flags to ensure high coverage contigs were not removed and that only duplications at the end of contigs were removed. The purged assembly and discarded sequences were then extracted.

### Mitochondrial Identification and Removal of Mitochondrial Contigs

MitoHiFi v3.2.1 (Uliano-Silva et al. 2023) was then used to query the purged assembly against the existing *Python regius* mtDNA genome (GenBank: AB177878.1, Dong and Kumazawa 2005) and identify mitochondrial contigs. Identified contigs were removed from the assembly using SeqKit v0.9.0 (Shen et al. 2016). QUAST/BUSCO analysis of the cleaned assembly was conducted as above (see *Genome quality control*).

### Repeat modeling and masking

RepeatModeler v2.0.6 (Flynn et al. 2020) was used to identify and model repeats in the cleaned assembly. The repeat family library from this *de novo* identification was merged with the sauropsida library from the Dfam database v3.8. RepeatMasker v4.1.7 (Smit, A et al. 2013) was used to softmask the repeats contained within the merged library, generate repeat statistics, and create a GFF repeat annotation file. The final cleaned and repeat-masked assembly was sorted by contig size.

### RNA Extraction and Sequencing

Frozen tissues (cardiac ventricle, liver, skeletal muscle, small intestine, brain, kidney, cardiac adipose, visceral adipose, ovary) were homogenized in Trizol with a mechanical homogenizer (Omni International) and incubated for 5 minutes at room temperature. Chloroform was added to the tubes at 1:5 vol/vol ratio and the samples were shaken, before incubating for 15 minutes at room temperature. Samples were subsequently centrifuged (12,000 RCF, 15 minutes, 4°C) and the aqueous upper layer was collected and combined with an equal volume of 75% ethanol and briefly vortexed. The solution was then applied to a RNeasy mini column (Qiagen) to help remove impurities. The manufacturer’s protocol was followed to wash the column in subsequent steps and then the RNA was eluted in RNase-free water. RNA samples were sent to Novogene Corporation (Sacramento, CA) for library prep and short-read Illumina NovaSeq sequencing at a minimum 50 million read-pair depth per sample. For long-read Nanopore sequencing, an RNA library was prepared with the PCR cDNA Barcoding kit (SQK-PCB111.24, Oxford Nanopore Technologies) and sequenced on an R9.4.1 PromethION flow cell (Oxford Nanopore Technologies).

### RNA read processing and alignment

Dorado v0.8.2 (Oxford Nanopore Technologies) was used to basecall Nanopore reads with the super high accuracy model and the –no-trim flag. Basecalled reads were then demultiplexed with Dorado and the --emit-fastq --no-trim flags provided. Pychopper v2.7.10 (Oxford Nanopore Technologies) was used to polish and trim the cDNA reads. The high-confidence full length reads and rescued reads were concatenated and mapped to the assembled genome with Minimap2 v2.17 with the splice preset (-x splice). Illumina reads were trimmed with bbduk.sh and the trimming parameters as for the Illumina WGS reads (see above). A Hisat2 index was constructed with Hisat2 v2.1.0 (Kim et al. 2019) and trimmed Illumina reads were aligned to the final genome assembly with sensitive alignment parameters and the --dta flag to generate alignments suitable for transcript assembly. SAM files containing Minimap2 and Hisat2 alignments were converted to BAM files, sorted, and indexed with Samtools. The 9 BAM files containing alignments from the Illumina reads (1 per tissue) were merged with the Samtools merge function.

### Genome annotation

The BRAKER3 annotation pipeline (Gabriel et al. 2024) was used to generate high confidence gene predictions. BRAKER3 incorporates RNA-seq and protein data and uses the gene prediction tools GeneMark-ETP (Brůna et al. 2024) and AUGUSTUS (Stanke et al. 2008) to train and predict genes with high support from extrinsic evidence. Final predictions from both GeneMark and AUGUSTUS are combined using TSEBRA (Gabriel et al. 2021) to produce the final annotation output. BRAKER3 was run with aligned RNA-Seq reads (BAM files) as extrinsic evidence and the pre-partitioned vertebrate set of proteins from the OrthoDB v.12 database (Kriventseva et al. 2015).

## Results and Discussion

### Assembly

We generated 51.5 Gb of Nanopore ONT reads with a mean length of 3847.6 bp and a read length N50 of 6377 bp (Table 1). Using the published Burmese python (*Python bivittatus*) genome size of 1.44 Gb (Castoe et al. 2013) this gave us an initial estimate of 35.8x coverage. The initial Flye assembly consisted of 3056 contigs with a reported mean coverage of 28x. Pilon was used to polish the genome assembly with aligned Illumina reads, which had a mean coverage of 56x (Table 2). We then cleaned the genome by removing all contigs smaller than 3000 bp, low coverage contigs, and contigs designated as haplotigs or collapsed repeats. We detected no evidence of contamination from bacteria or other organisms.

**Table 1.** Nanopore DNA sequencing read statistics.

|  |  |
| --- | --- |
| **Mean read length (bp)** | 3847.6 |
| **Mean read quality** | 11.3 |
| **Median read length (bp)** | 2221.0 |
| **Median read quality** | 18.3 |
| **# of Reads** | 13,378,060 |
| **Read Length N50** | 6377 |
| **STDEV read length** | 7558.5 |
| **Total bases** | 51,473,874,864 |

**Table 2.** Illumina NovaSeq WGS read statistics.

|  | **R1** | **R2** |
| --- | --- | --- |
| **# of Reads** | 419,719,743 | 419,719,743 |
| **Mean Sequence Quality** | 38.19 | 37.76 |
| **# Trimmed Reads** | 418,142,934 | 418,142,934 |
| **Mean Sequence Quality – Trimmed Reads** | 38.34 | 38.08 |

The final genome assembly was 1.45 Gb, virtually identical in size to the genomes of the closely related Burmese python and Reticulated python (*Malayopython reticulatus*), which are both approximately 1.44 Gb (De Smet 1981; Castoe et al. 2013). Unsurprisingly, our assembly was also significantly more contiguous than the Burmese genome which was sequenced over a decade ago via shotgun sequencing. The final polished and purged assembly consisted of 1,077 contigs, approximately 36 times less than the 39,112 contigs of the Burmese python assembly (Table 3). The contig N50 value was significantly higher for the assembled ball python genome (18.1 Mb) compared to that of the published Burmese python genome (10.7 Kb). It should be noted that the Burmese python genome was assembled to the scaffold level yet still is much less contiguous than our contig level assembly. The final assembly has a BUSCO score of 98% (vertebrate complete and single-copy orthologs, Table 4), noticeably higher than 89.7% for the Burmese python genome. The reduced BUSCO score for the Burmese genome is due to a higher percentage of fragmented and missing orthologs compared to our assembly (Table 4). Furthermore, our genome is slightly higher than the 91.5-96.1% range reported in a study that assembled and analyzed 14 *de novo* genomes from 12 snake families (Peng et al. 2023). Consequently, our assembly represents the highest quality and most contiguous genome for a member of the *Pythonidae* family to date, and is line with the quality of recently published snake genomes (Peng et al. 2023; Westeen et al. 2023).

**Table 3.**
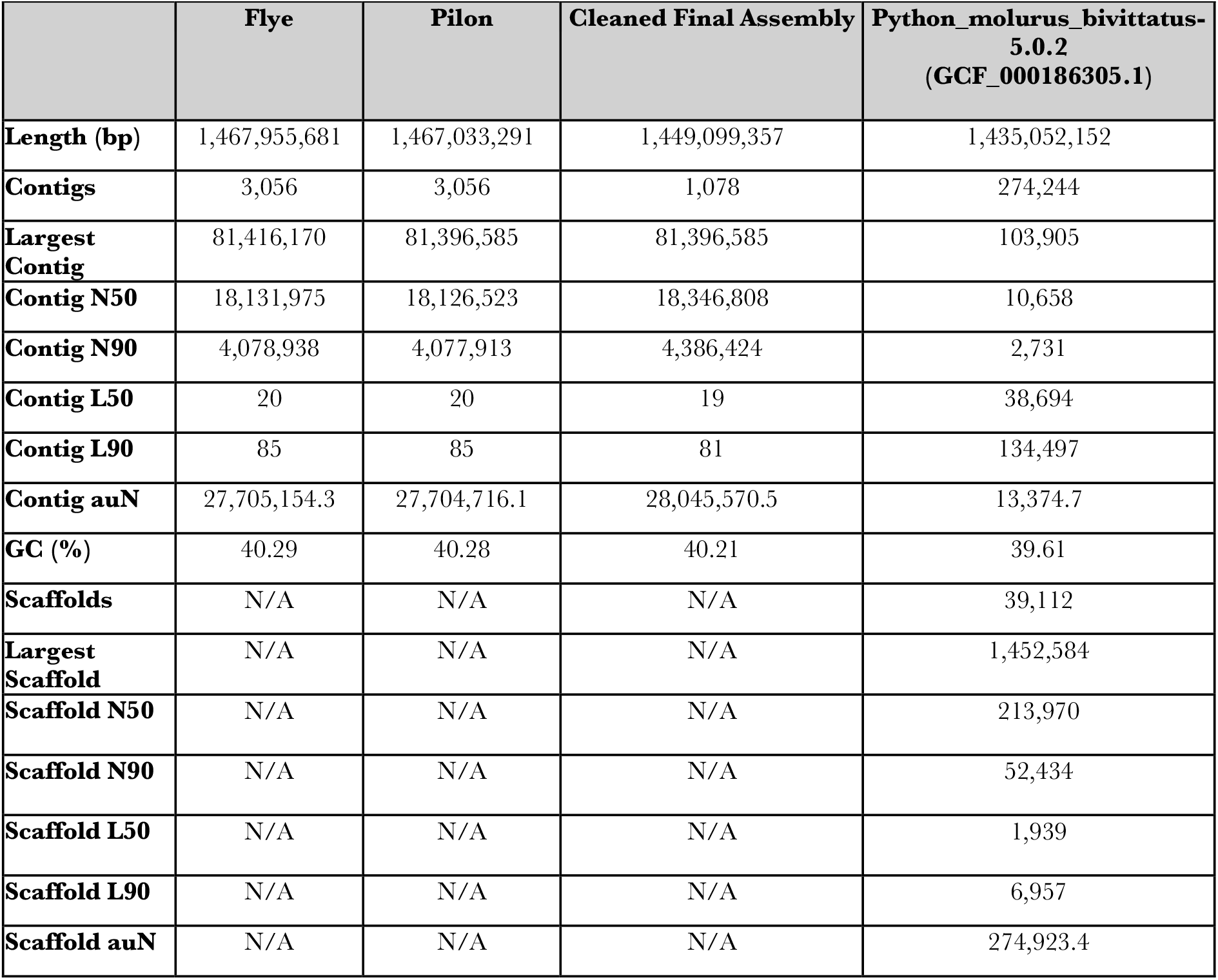
Quality statistics for the Flye, Pilon-polished, and final cleaned *P. regius* genome assemblies. The statistics for the latest NCBI RefSeq version of the *P. bivittatus* genome are provided for comparison.

**Table 4.**
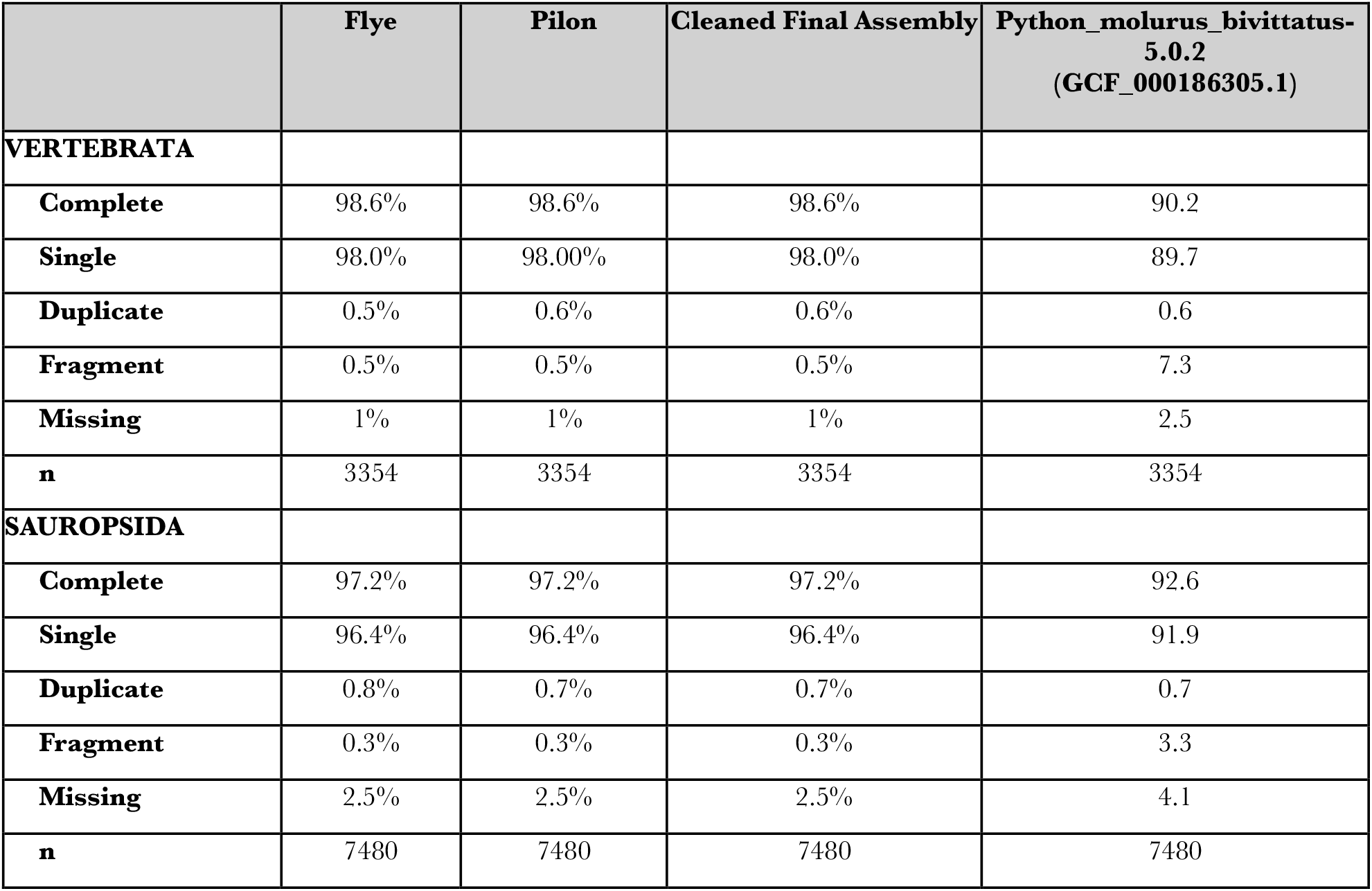
Genome BUSCO scores for the Flye, Pilon-polished, and final cleaned *P. regius* assemblies. The statistics for the latest NCBI RefSeq version of the *P. bivittatus* genome are provided for comparison. Both the vertebrate_odb10 and sauropsida_odb10 databases were used for BUSCO analysis.

### Repeat annotation

The repeat element content of snake genomes can vary significantly, with reported ranges from 25-73% (Ahmad et al. 2021). We masked 34.8% of the final ball python assembly, similar to the 31.8 % and 35.2% values for the Burmese python and King Cobra respectively (Castoe et al. 2013) (Table 5). Recent evidence suggests a correlation between genome size and repeat content in snakes (Peng et al. 2023). Consistent with this, the repeat content and genome size for ball pythons is on the lower end among other snake species. Additionally, CR1 and LTR elements have been found to be less abundant in primitive snakes such as pythons and boas compared to more advanced snake species (Castoe et al. 2011; Yin et al. 2016; Ahmad et al. 2021). Our data is consistent with this finding when comparing the repeat contents between the ball python genome and the genomes of recently sequenced and repeat-annotated snake genomes (Ahmad et al. 2021; Peng et al. 2023). Interestingly, ball pythons have a higher long interspersed nuclear element (LINE) content (13.4%) than Burmese pythons (8.6%). Whether this biological or a consequence of the poor contiguity of the existing Burmese python genome is unclear.

**Table 5.**
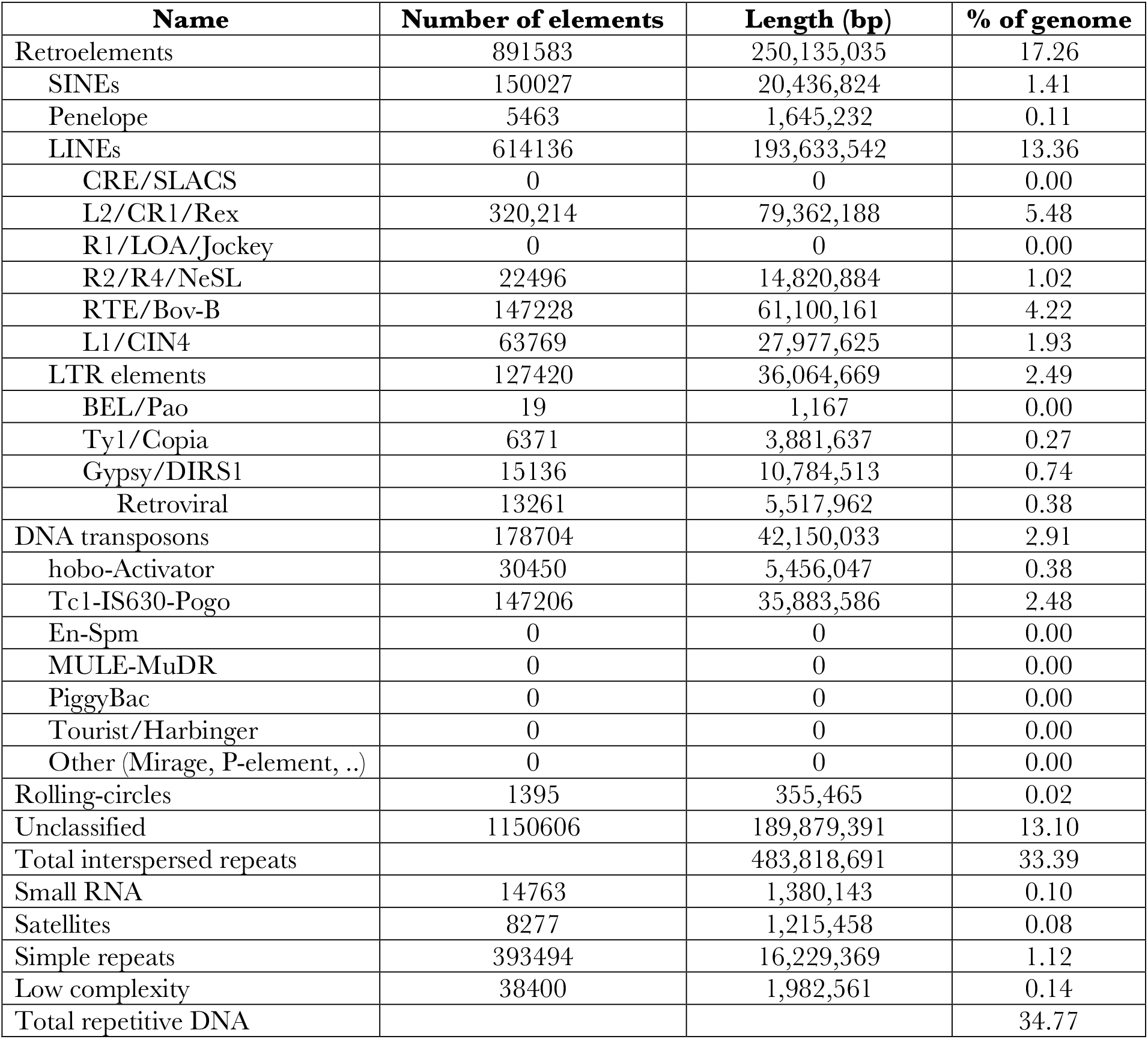
Repetitive DNA content of the *P. regius* genome.

| **Name** | **Number of elements** | **Length (bp)** | **% of genome** |
| --- | --- | --- | --- |
| Retroelements | 891583 | 250,135,035 | 17.26 |
| SINEs | 150027 | 20,436,824 | 1.41 |
| Penelope | 5463 | 1,645,232 | 0.11 |
| LINEs | 614136 | 193,633,542 | 13.36 |
| CRE/SLACS | 0 | 0 | 0.00 |
| L2/CR1/Rex | 320,214 | 79,362,188 | 5.48 |
| R1/LOA/Jockey | 0 | 0 | 0.00 |
| R2/R4/NeSL | 22496 | 14,820,884 | 1.02 |
| RTE/Bov-B | 147228 | 61,100,161 | 4.22 |
| L1/CIN4 | 63769 | 27,977,625 | 1.93 |
| LTR elements | 127420 | 36,064,669 | 2.49 |
| BEL/Pao | 19 | 1,167 | 0.00 |
| Ty1/Copia | 6371 | 3,881,637 | 0.27 |
| Gypsy/DIRS1 | 15136 | 10,784,513 | 0.74 |
| Retroviral | 13261 | 5,517,962 | 0.38 |
| DNA transposons | 178704 | 42,150,033 | 2.91 |
| hobo-Activator | 30450 | 5,456,047 | 0.38 |
| Tc1-IS630-Pogo | 147206 | 35,883,586 | 2.48 |
| En-Spm | 0 | 0 | 0.00 |
| MULE-MuDR | 0 | 0 | 0.00 |
| PiggyBac | 0 | 0 | 0.00 |
| Tourist/Harbinger | 0 | 0 | 0.00 |
| Other (Mirage, P-element, ..) | 0 | 0 | 0.00 |
| Rolling-circles | 1395 | 355,465 | 0.02 |
| Unclassified | 1150606 | 189,879,391 | 13.10 |
| Total interspersed repeats |  | 483,818,691 | 33.39 |
| Small RNA | 14763 | 1,380,143 | 0.10 |
| Satellites | 8277 | 1,215,458 | 0.08 |
| Simple repeats | 393494 | 16,229,369 | 1.12 |
| Low complexity | 38400 | 1,982,561 | 0.14 |
| Total repetitive DNA |  |  | 34.77 |

### Genome annotation

RNA from nine different tissues was extracted and sequenced via Illumina and Nanopore cDNA sequencing. Across the nine tissues we generated 37,164,799 Nanopore reads with an average read length N50 of 705.2 bp, as well as over 592 million high-quality Illumina read pairs (Table 6, Table 7). We identified 18,842 putative coding genes via the BRAKER3 genome annotation pipeline (Table 8). BUSCO analysis of the longest proteins isoforms from these genes yielded scores of 97.1% (95.9% single copy and 1.2% duplicates) against the vertebrate database and 93.7% (92.4% single copy and 1.3% duplicates) against the sauropsida database (Table 9).

**Table 6.** Nanopore cDNA sequencing statistics used for genome annotation.

|  | Ovary | Ventricle | Liver | Skeletal Muscle | Small Intestine | Brain | Kidney | Cardiac Adipose | Visceral Adipose |
| --- | --- | --- | --- | --- | --- | --- | --- | --- | --- |
| Mean read length (bp) | 841.1 | 449.9 | 342.5 | 884.9 | 483.3 | 570.7 | 346.6 | 631.9 | 622.0 |
| Mean read quality | 10.4 | 9.7 | 9.0 | 10.1 | 9.9 | 10.1 | 9.3 | 9.9 | 9.9 |
| Median read length (bp) | 747.0 | 292.0 | 258.0 | 906.0 | 411.0 | 434.0 | 268.0 | 568.0 | 515.0 |
| Median read quality | 11.9 | 10.8 | 10.1 | 11.7 | 11.1 | 11.5 | 10.5 | 11.2 | 11.3 |
| # of Reads | 1,444,246 | 5,307,918 | 7,938,877 | 1,641,433 | 3,534,382 | 3,675,177 | 8,007,701 | 3,124,097 | 2,490,968.0 |
| Read Length N50 | 1,039.0 | 606.0 | 319.0 | 1,149.0 | 613.0 | 736.0 | 331.0 | 780.0 | 774.0 |
| STDEV read length | 531.0 | 348.4 | 303.3 | 529.5 | 348.7 | 453.0 | 332.7 | 475.9 | 445.7 |
| Total bases | 1,215,144,379 | 2,388,039,699 | 2,719,437,803 | 1,452,439,846 | 1,708,134,062 | 2,097,289,662 | 2,775,554,268 | 1,974,226,081 | 1,549,318,650 |

**Table 7.** Illumina NovaSeq RNA sequencing statistics used for genome annotation.

| Tissue | Read Pairs |  | R1 Mean Sequence Quality |  | R2 Mean Sequence Quality |  |
| --- | --- | --- | --- | --- | --- | --- |
|  | Raw | Trimmed | Raw | Trimmed | Raw | Trimmed |
| Ovary | 64,956,172 | 64,955,137 | 38.55 | 38.61 | 38.48 | 38.64 |
| Ventricle | 75,089,332 | 75,087,842 | 38.61 | 38.68 | 38.53 | 38.73 |
| Liver | 100,579,525 | 100,578,013 | 38.58 | 38.65 | 38.51 | 38.65 |
| Skeletal Muscle | 57,581,057 | 57,580,062 | 38.62 | 38.67 | 38.59 | 38.69 |
| Small Intestine | 58,340,546 | 58,339,660 | 38.59 | 38.64 | 38.55 | 38.64 |
| Brain | 56,945,386 | 56,944,575 | 38.51 | 38.58 | 38.40 | 38.53 |
| Kidney | 61,858,414 | 61,857,151 | 38.55 | 38.61 | 38.34 | 38.50 |
| Cardiac Adipose | 53,914,022 | 53,913,057 | 38.58 | 38.66 | 38.45 | 38.62 |
| Visceral Adipose | 62,870,966 | 62,870,017 | 38.67 | 38.71 | 38.64 | 38.70 |
| Total | 592,135,420 | 592,125,514 | 38.59 | 38.65 | 38.50 | 38.64 |

**Table 8.** Quantitative summary of the BRAKER3 annotation of the final *P. regius* genome.

| **Feature** | **Count** |
| --- | --- |
| Genes | 18,842 |
| Transcripts/Proteins | 25,522 |
| Mean gene length (bp) | 28,996 |
| Exons | 286,501 |
| Introns | 260,979 |
| Mean exons per transcript | 11.2 |
| Mean introns per transcript | 10.2 |
| Mean exon length (bp) | 164 |
| Mean intron length (bp) | 3168 |
| Single exon transcripts | 2029 |

**Table 9.** Proteome BUSCO Scores for both *P*.*regius* and *P. bivittatus*.

|  | **Python regius**<br>**(BRAKER3 annotation)** | **Python_molurus_bivittatus-**<br>**5.0.2**<br>**(GCF_000186305.1)** |
| --- | --- | --- |
| **VERTEBRATA** |  |  |
| **Complete** | 97.1 | 90.8 |
| **Single** | 95.9 | 89.7 |
| **Duplicate** | 1.2% | 1.1 |
| **Fragment** | 0.4% | 6.4 |
| **Missing** | 2.4% | 2.7 |
| **n** | 3354 | 3354 |
| **SAUROPSIDA** |  |  |
| **Complete** | 93.7% | 92.2 |
| **Single** | 92.4% | 91.2 |
| **Duplicate** | 1.3% | 1.0 |
| **Fragment** | 0.6% | 3.5 |
| **Missing** | 5.7% | 4.3 |
| **n** | 7480 | 7480 |

## Conclusion

We present the first genome assembly for the ball python, and the highest quality genome assembly to date for a member of the *Pythonidae* family. From a general standpoint, this work lays the foundation for further exploration of genetic diversity among snakes and broader evolutionary questions in the field of developmental biology. It will also aid in studies seeking to identify the molecular basis behind resistance to muscular atrophy, as well as the ability of pythons to undergo significant, rapid, and reversible cardiac remodeling following a meal. Such studies may provide unique insights into mechanisms of muscular atrophy and heart disease in mammals and open potential avenues for novel therapeutics targeted towards these conditions.

## Data Availability

Sequencing data, genome assembly, and genome annotation files will be uploaded to NCBI prior to submission. Data will be included under BioProject ID PRJNA1217506.

## Acknowledgements

We would like to thank the University of Colorado Boulder Office of Animal Resources for assistance with python colony husbandry and all those who assisted with python dissections, especially Jack Gugel and Skip Maas. We would also like to thank Yuxiao Tan for taking and providing the picture used in Figure 1.

## References

Ahmad SF, Singchat W, Panthum T, Srikulnath K. 2021. Impact of Repetitive DNA Elements on Snake Genome Biology and Evolution. Cells. 10(7):1707. 10.3390/cells10071707

Altschul SF, Gish W, Miller W, Myers EW, Lipman DJ. 1990. Basic local alignment search tool. J Mol Biol. 215(3):403–410. 10.1016/S0022-2836(05)80360-2

Andersen JB, Rourke BC, Caiozzo VJ, Bennett AF, Hicks JW. 2005. Physiology: postprandial cardiac hypertrophy in pythons. Nature. 434(7029):37–38. 10.1038/434037a

Brůna T, Lomsadze A, Borodovsky M. 2024. GeneMark-ETP significantly improves the accuracy of automatic annotation of large eukaryotic genomes. Genome Res. 34(5):757–768. 10.1101/gr.278373.123

Buchfink B, Reuter K, Drost H-G. 2021. Sensitive protein alignments at tree-of-life scale using DIAMOND. Nat Methods. 18(4):366–368. 10.1038/s41592-021-01101-x

Bushnell B. BBMap. BBMap [Internet]. [accessed 2025 Jan 27]. https://sourceforge.net/projects/bbmap/

Camacho C, Coulouris G, Avagyan V, Ma N, Papadopoulos J, Bealer K, Madden TL. 2009. BLAST+: architecture and applications. BMC Bioinformatics. 10(1):421. 10.1186/1471-2105-10-421

Castoe TA, Hall KT, Guibotsy Mboulas ML, Gu W, de Koning APJ, Fox SE, Poole AW, Vemulapalli V, Daza JM, Mockler T, et al. 2011. Discovery of Highly Divergent Repeat Landscapes in Snake Genomes Using High-Throughput Sequencing. Genome Biol Evol. 3:641–653. 10.1093/gbe/evr043

Castoe TA, Jiang ZJ, Gu W, Wang ZO, Pollock DD. 2008. Adaptive Evolution and Functional Redesign of Core Metabolic Proteins in Snakes. PLOS ONE. 3(5):e2201. 10.1371/journal.pone.0002201

Castoe TA, de Koning APJ, Hall KT, Card DC, Schield DR, Fujita MK, Ruggiero RP, Degner JF, Daza JM, Gu W, et al. 2013. The Burmese python genome reveals the molecular basis for extreme adaptation in snakes. Proceedings of the National Academy of Sciences. 110(51):20645–20650. 10.1073/pnas.1314475110

Challis R, Richards E, Rajan J, Cochrane G, Blaxter M. 2020. BlobToolKit – Interactive Quality Assessment of Genome Assemblies. G3 Genes|Genomes|Genetics. 10(4):1361–1374. 10.1534/g3.119.400908

Crocini C, Woulfe KC, Ozeroff CD, Perni S, Cardiello J, Walker CJ, Wilson CE, Anseth K, Allen MA, Leinwand LA. 2024. Postprandial cardiac hypertrophy is sustained by mechanics, epigenetic, and metabolic reprogramming in pythons. Proceedings of the National Academy of Sciences. 121(36):e2322726121. 10.1073/pnas.2322726121

Danecek P, Bonfield JK, Liddle J, Marshall J, Ohan V, Pollard MO, Whitwham A, Keane T, McCarthy SA, Davies RM, Li H. 2021. Twelve years of SAMtools and BCFtools. GigaScience. 10(2):giab008. 10.1093/gigascience/giab008

De Coster W, D’Hert S, Schultz DT, Cruts M, Van Broeckhoven C. 2018. NanoPack: visualizing and processing long-read sequencing data. Bioinformatics. 34(15):2666–2669. 10.1093/bioinformatics/bty149

De Smet W. 1981. The nuclear Feulgen-DNA content of the vertebrates (especially reptiles), as measured by fluorescence cytophotometry, with notes on the cell and chromosome size. Acta Zool et Pathologica Antverpiensia. 76(1):119–167.

Dong S, Kumazawa Y. 2005. Complete mitochondrial DNA sequences of six snakes: phylogenetic relationships and molecular evolution of genomic features. J Mol Evol. 61(1):12–22. 10.1007/s00239-004-0190-9

Flynn JM, Hubley R, Goubert C, Rosen J, Clark AG, Feschotte C, Smit AF. 2020. RepeatModeler2 for automated genomic discovery of transposable element families. Proceedings of the National Academy of Sciences. 117(17):9451–9457. 10.1073/pnas.1921046117

Gabriel L, Brůna T, Hoff KJ, Ebel M, Lomsadze A, Borodovsky M, Stanke M. 2024. BRAKER3: Fully automated genome annotation using RNA-seq and protein evidence with GeneMark-ETP, AUGUSTUS, and TSEBRA. Genome Res. 34(5):769–777. 10.1101/gr.278090.123

Gabriel L, Hoff KJ, Brůna T, Borodovsky M, Stanke M. 2021. TSEBRA: transcript selector for BRAKER. BMC Bioinformatics. 22(1):566. 10.1186/s12859-021-04482-0

Guan D, McCarthy SA, Wood J, Howe K, Wang Y, Durbin R. 2020. Identifying and removing haplotypic duplication in primary genome assemblies. Bioinformatics. 36(9):2896–2898. 10.1093/bioinformatics/btaa025

Kim D, Paggi JM, Park C, Bennett C, Salzberg SL. 2019. Graph-based genome alignment and genotyping with HISAT2 and HISAT-genotype. Nat Biotechnol. 37(8):907–915. 10.1038/s41587-019-0201-4

Kolmogorov M, Yuan J, Lin Y, Pevzner PA. 2019. Assembly of long, error-prone reads using repeat graphs. Nat Biotechnol. 37(5):540–546. 10.1038/s41587-019-0072-8

Kriventseva EV, Tegenfeldt F, Petty TJ, Waterhouse RM, Simão FA, Pozdnyakov IA, Ioannidis P, Zdobnov EM. 2015. OrthoDB v8: update of the hierarchical catalog of orthologs and the underlying free software. Nucleic Acids Res. 43(Database issue):D250–256. 10.1093/nar/gku1220

Li H. 2013. Aligning sequence reads, clone sequences and assembly contigs with BWA-MEM [Internet]. [accessed 2025 Jan 27]. 10.48550/arXiv.1303.3997

Li H. 2018. Minimap2: pairwise alignment for nucleotide sequences. Bioinformatics. 34(18):3094–3100. 10.1093/bioinformatics/bty191

Li H, Handsaker B, Wysoker A, Fennell T, Ruan J, Homer N, Marth G, Abecasis G, Durbin R, 1000 Genome Project Data Processing Subgroup. 2009. The Sequence Alignment/Map format and SAMtools. Bioinformatics. 25(16):2078–2079. 10.1093/bioinformatics/btp352

Manni M, Berkeley MR, Seppey M, Simão FA, Zdobnov EM. 2021. BUSCO Update: Novel and Streamlined Workflows along with Broader and Deeper Phylogenetic Coverage for Scoring of Eukaryotic, Prokaryotic, and Viral Genomes. Molecular Biology and Evolution. 38(10):4647–4654. 10.1093/molbev/msab199

Martin TG, Hunt DR, Langer SJ, Tan Y, Ebmeier CC, Leinwand LA. 2024. Regression of postprandial cardiac hypertrophy in burmese pythons is mediated by FoxO1. Proceedings of the National Academy of Sciences. 121(41):e2408719121. 10.1073/pnas.2408719121

McCue MD. 2007. Snakes survive starvation by employing supply- and demand-side economic strategies. Zoology (Jena). 110(4):318–327. 10.1016/j.zool.2007.02.004

Mikheenko A, Saveliev V, Hirsch P, Gurevich A. 2023. WebQUAST: online evaluation of genome assemblies. Nucleic Acids Research. 51(W1):W601–W606. 10.1093/nar/gkad406

Peng C, Wu D-D, Ren J-L, Peng Z-L, Ma Z, Wu W, Lv Y, Wang Z, Deng C, Jiang K, et al. 2023. Large-scale snake genome analyses provide insights into vertebrate development. Cell. 186(14):2959-2976.e22. 10.1016/j.cell.2023.05.030

Riquelme CA, Magida JA, Harrison BC, Wall CE, Marr TG, Secor SM, Leinwand LA. 2011. Fatty acids identified in the Burmese python promote beneficial cardiac growth. Science. 334(6055):528–531. 10.1126/science.1210558

Secor SM, Diamond J. 1995. Adaptive responses to feeding in Burmese pythons: pay before pumping. J Exp Biol. 198(Pt 6):1313–1325. 10.1242/jeb.198.6.1313

Secor SM, Diamond J. 1998. A vertebrate model of extreme physiological regulation. Nature. 395(6703):659–662. 10.1038/27131

Shen W, Le S, Li Y, Hu F. 2016. SeqKit: A Cross-Platform and Ultrafast Toolkit for FASTA/Q File Manipulation. PLOS ONE. 11(10):e0163962. 10.1371/journal.pone.0163962

Smit, A, Hubley, R, Green, P. 2013. RepeatMasker Open-4.0 [Internet]. http://www.repeatmasker.org

Stanke M, Diekhans M, Baertsch R, Haussler D. 2008. Using native and syntenically mapped cDNA alignments to improve de novo gene finding. Bioinformatics. 24(5):637–644. 10.1093/bioinformatics/btn013

Stenvinkel P, Painer J, Johnson RJ, Natterson-Horowitz B. 2020. Biomimetics - Nature’s roadmap to insights and solutions for burden of lifestyle diseases. J Intern Med. 287(3):238–251. 10.1111/joim.12982

Tan Y, Martin TG, Harrison BC, Leinwand LA. 2023. Utility of the burmese Python as a model for studying plasticity of extreme physiological systems. J Muscle Res Cell Motil. 44(2):95–106. 10.1007/s10974-022-09632-2

Uliano-Silva M, Ferreira JGRN, Krasheninnikova K, Blaxter M, Mieszkowska N, Hall N, Holland P, Durbin R, Richards T, Kersey P, et al. 2023. MitoHiFi: a python pipeline for mitochondrial genome assembly from PacBio high fidelity reads. BMC Bioinformatics. 24(1):288. 10.1186/s12859-023-05385-y

Vonk FJ, Richardson MK. 2008. Serpent clocks tick faster. Nature. 454(7202):282–283. 10.1038/454282a

Walker BJ, Abeel T, Shea T, Priest M, Abouelliel A, Sakthikumar S, Cuomo CA, Zeng Q, Wortman J, Young SK, Earl AM. 2014. Pilon: An Integrated Tool for Comprehensive Microbial Variant Detection and Genome Assembly Improvement. PLOS ONE. 9(11):e112963. 10.1371/journal.pone.0112963

Westeen EP, Escalona M, Beraut E, Marimuthu MPA, Nguyen O, Fisher RN, Toffelmier E, Shaffer HB, Wang IJ. 2023. A reference genome assembly for the continentally distributed ring-necked snake, Diadophis punctatus. Journal of Heredity. 114(6):690–697. 10.1093/jhered/esad051

Whibley A, Kelley JL, Narum SR. 2021. The changing face of genome assemblies: Guidance on achieving high-quality reference genomes. Molecular Ecology Resources. 21(3):641–652. 10.1111/1755-0998.13312

Yin W, Wang Z, Li Q, Lian J, Zhou Y, Lu B, Jin L, Qiu P, Zhang P, Zhu W, et al. 2016. Evolutionary trajectories of snake genes and genomes revealed by comparative analyses of five-pacer viper. Nat Commun. 7:13107. 10.1038/ncomms13107

